# Ancestrally Shared Regenerative Mechanisms Across the Metazoa: A Transcriptomic Case Study in the Demosponge *Halisarca caerulea*

**DOI:** 10.1101/160689

**Authors:** Nathan J. Kenny, Jasper M. de Goeij, Didier M. de Bakker, Casey G. Whalen, Eugene Berezikov, Ana Riesgo

**Affiliations:** Life Sciences Department, The Natural History Museum, Cromwell Road, London SW7 5BD, UK; Department of Freshwater and Marine Science, University of Amsterdam, Postbus 94240, 1090 GE Amsterdam, The Netherlands; European Research Institute for the Biology of Ageing, University of Groningen, University Medical Center Groningen, Groningen, The Netherlands

**Keywords:** Regeneration, transcriptome, *Halisarca caerulea*, Porifera, ancestral cassette

## Abstract

Regeneration is an essential process for all multicellular organisms, allowing them to recover effectively from internal and external injury. This process has been studied extensively in a medical context in vertebrates, with pathways often investigated mechanistically, both to derive increased understanding and as potential drug targets for therapy. Several species from other parts of the metazoan tree of life, noted for their regenerative prowess, have previously been targeted for study. This allows us to understand their regenerative mechanisms and see how they could be adapted for use in medicine. Species in clades such as *Hydra,* planarians and echinoderms can regenerate large portions of their body, the former two clades being able to completely regenerate from even a small portion of their somatic tissue. Less well-documented for their regenerative abilities are sponges. This is surprising, as they are both one of the earliest-branching extant metazoan phyla on Earth, and are rapidly able to respond to injury. Their sessile lifestyle, lack of an external protective layer, inability to respond to predation and filter-feeding strategy all mean that regeneration is often required. In particular the demosponge genus *Halisarca* has been noted for its fast cell turnover and ability to quickly adjust its cell kinetic properties to repair damage through regeneration. However, while the rate and structure of regeneration in sponges has begun to be investigated, the molecular mechanisms behind this ability are yet to be catalogued.

Here we describe the assembly of a reference transcriptome for *Halisarca caerulea,* along with additional transcriptomes noting response to injury before, shortly following (2 hrs post-), and 12 hrs after trauma. RNAseq reads were assembled using Trinity, annotated, and samples compared, to allow initial insight into the transcriptomic basis of sponge regenerative processes. These resources are deep, with our reference assembly containing more than 92.6% of the BUSCO Metazoa set of genes, and well-assembled (N50s of 836, 957, 1,688 and 2,032 for untreated, 2h, 12h and reference transcriptomes respectively), and therefore represent excellent initial resources as a bedrock for future study. The generation of transcriptomic resources from sponges before and following deliberate damage has allowed us to study particular pathways within this species responsible for repairing damage. We note particularly the involvement of the Wnt cascades in this process in this species, and detail the contents of this cascade, along with cell cycle, extracellular matrix and apoptosis-linked genes in this work.

This resource represents an excellent starting point for the continued development of this knowledge, given *H. caerulea*’s ability to regenerate and position as an outgroup for comparing the process of regeneration across metazoan lineages. With this resource in place, we can begin to infer the regenerative capacity of the common ancestor of all extant animal life, and unravel the elements of regeneration in an often-overlooked clade.

## Introduction

In order to survive the often-hostile conditions in which they live, multicellular organisms have evolved the ability to regenerate their tissues, allowing them to recover if injured. While almost all multicellular organisms can regenerate throughout their lives (Sánchez Alvarado 2000), some clades are particularly adept at this process, and have been subjected to particular scrutiny. Sponges are well known for their regeneration abilities (Wulff 2010), capable of regenerating a lost fragment, regrowing from a small piece, or even regenerating entirely from disaggregated cells (Korotkova 1997). They are capable of regenerating at rates of as much as 2900 times the normal growth rates (Ayling 1983) although this rate is by no means universal (Kahn and Leys 2016) and comes at the expense of standard cell cycling (Henry & Hart, 2005). This is perhaps a response to their niche, as they are benthic, non-motile organisms, which can be exposed to damage in a variety of ways. Their inability to avoid physical damage from predation is obvious, but their filter feeding lifestyle also exposes them to bacterial, viral, chemical and physical stresses, and they have no external protection from their environments (Wulff 2010). To date, studies of the ways sponges regenerate have tended to be histological (e.g. Korotkova, 1961; Korotkova, 1970; Korotkova, 1972; Korotkova and Movchan 1973; Thiney 1972; Boury-Esnault 1976). Only recently have these traditional studies, based on light microscopy, been followed by studies using electron microscopy (Borisenko et al. 2015, Coutinho et al 2017) and cell cycle kinetics (de Goeij et al. 2009, Alexander et al. 2015). While we already have a deeper understanding of the drivers of these processes in many phyla, in poriferans we still have little to no understanding of the molecular mechanisms underlying these regenerative processes. The advent of next-generation sequencing techniques allows us to solve this knowledge gap efficiently and effectively, and bring sponges into consideration as model organisms for understanding regeneration, given their peculiar advantages as described above.

While described only relatively recently, the demosponge *Halisarca caerulea* (Vacelet & Donadey, 1987) has become the subject of scientific interest due to its ability to recover from damage at a rapid speed (Alexander et al. 2015) and its ecological significance as key recycler of resources in tropical coral ecosystems (De Goeij et al. 2008, 2013). The cell cycle of *Halisarca caerulea* (and particularly that of choanocytes, the sponges filter cells) is exceptionally rapid, approximately 6 h, resulting in high cell turnover driven by choanocyte proliferation, followed by cell shedding (rather than apoptosis, de Goeij et al. 2009), and this may allow it to recover from external damage in an exceptionally efficient manner. The regeneration abilities of its congeneric species have been studied more extensively (e.g., Korotkova and Movchan 1973; Borisenko et al. 2015), where it was noted that the regeneration process is comprised by three steps: formation of regenerative plug (mucus, collagen, bacteria, debris and several cell types aggregate into this plug), reorganization of the extracellular matrix, and wound healing by blastema formation (transdifferentiation of archaeocytes into exopinacocytes). ‘Old’, shed cells (i.e. detritus) then form the basis of a sponge-mediated food chain named ‘the sponge loop’, in which sponge-derived detritus serves as one of the major recycled food sources on shallow water and deep-sea coral reefs (De Goeij et al. 2013; Rix et al. 2016).

Regeneration has been studied in a number of taxa, and we now know that it can occur in a variety of ways (Tanaka & Reddien, 2011; Vervoort, 2011, Tiozzo and Copley 2015). Although some aspects of its mechanics seem to be shared ancestrally, we have only begun to fully catalogue the molecular and cellular architecture of this process across the tree of life. Several organisms, known to be effective regenerators, have already been the focus of studies which have revealed that the a wide range of pathways, including FGF, Wnt and TGFβ pathways are common components of such a response (Sánchez Alvarado and Tsonis 2006). This knowledge, reinforced by unbiased screens looking for any genes involved in regeneration (e.g., Reddien et al. 2005) have revealed a variety of potential target genes involved in regenerative responses and allowed these pathways to be targeted for potential clinical use. However, we still have little idea as to which of these pathways are shared ancestrally across the tree of life (Tiozzo and Copley 2015), a key consideration when trying to apply findings from other Superphyla to human studies, as we look for effective treatments that are likely to be shared. Generally, the first molecular signals of regeneration in any metazoan phyla involve wound healing responses, led by Ca^2+^, hydrogen peroxide (H_2_O_2_) and ATP (Cordeiro and Jacinto, 2013, Tiozzo and Copley 2015), and these wound-healing responses are thought to have been present in the metazoan common ancestor. These are followed by gene-level responses to allow regeneration itself, which can occur in any of three ways (1) via extant stem cells, (2) through trans/de-differentiation of adult tissue, or (3), through proliferation of already extant tissue. Different species and phyla exhibit different combinations of these responses, and some of them seem to share the molecular mechanisms behind the responses themselves.

Sponges possess large populations of totipotent (archaeocyte) and pluripotent (choanocyte) stem cells present in adult tissue (Simpson 1984; Müller 2006, Funayama 2010, Funayama 2013). This means that regeneration could occur without complete de-differentiation or trans-differentiation. The main processes observed so far in regenerating tissue are epithelial to mesenchymal transition, dedifferentiation, cell migration, and blastema formation (e.g., Borisenko et al. 2015). It is possible that regeneration is a simpler process transcriptionally in these organisms, without the necessity for full genetic reprogramming before it begins to occur (Tiozzo and Copley 2015). This could make sponges a potent source of information regarding the basic signals involved in beginning a regenerative response, and it is known that sponges, urochordates and flatworms share the use of the molecular markers *piwi* and *nanos* in stem cell niche maintenance (Brown et al. 2009, de Mulder et al. 2009, Funayama 2013). Further signals of stem cell response to regenerative need could be discerned by comparing the transcriptional responses of these disparate phyla to injury.

As sponges are among the earliest branching phyla, if not the earliest branching metazoan phylum (Halanych 2015, Simion et al 2017), the study of regeneration in these species will be of widespread benefit. Comparative methods, with sponges as an outgroup, will allow us to understand the evolution of regeneration in specific clades, and know which traits are likely to be ancestrally shared. Furthermore, the simple structure and tissues of sponges make them excellent models for understanding the very fundamental aspects of wound healing (e.g., Alexander et al. 2015). As aforementioned, sponges are increasingly recognized as key ecosystem drivers and engineers in both shallow and deep-sea reefs (De Goeij et al. 2013, Rix et al. 2016), which is hypothesized to be driven by their fast cell turnover (De Goeij et al. 2009, 2013; Alexander et al. 2014). As a result of this, sponges also can heal much faster than some animals, simplifying experimental procedures.

Our efforts to understand sponge regeneration in particular and biology more generally are still hampered by depauperate sequence resources, and genomic and transcriptomic sequences in this clade remain limited in number. Within the Porifera, few species have been studied at the genomic level: *Amphimedon queenslandica* (Srivistava et al. 2010) is the most complete published example, and *Sycon ciliatum* (Leininger et al. 2014), *Oscarella carmela* (Nichols et al. 2012) and *Tethya wilhelma* (Francis et al. 2017) have been sequenced to a draft level. Several studies of sponges have also been completed using transcriptomic sequencing, including in the related sponge species *Halisarca dujardini,* but these are from complete adult and larval samples, and are unlinked to regeneration (Borisenko et al. 2016). To help remedy this, and to provide the first insights into the molecular means of regeneration in the Porifera, we have undertaken a study describing transcriptional changes in the demosponge *H. caerulea* before, 2 hours after, and 12 hours following injury. This study represents, to our knowledge, the first examining regeneration in sponges using NGS techniques. We have provided a deep transcriptomic resource describing the cassette of activated genes before, during and following an induced regenerational response. This has allowed us to describe some of the most influential pathways involved in moderating such responses. This study will be useful in understanding regeneration in sponges and more broadly across the tree of metazoan life, and its timeliness and depth will provide a basis for a range of further studies, building on the initial observations made in this manuscript.

## Material and Methods

### Tissue Collection, Sample Treatments and RNA Extraction

Sponges were collected by SCUBA diving at water depths of 15–25 m near the CARMABI Foundation Research Station in Curaçao (Lat/Long: 12°12′N, 68°56′W) in February–April 2013. This work was in compliance with a research permit issued by the Curaçaoan Ministry of Health (#2012/48584). Specimens of the thin (± 2 mm) encrusting sponge *H. caerulea* were collected by chiselling sponge specimens (± 25 cm^2^) and their immediate coral rock substratum from the larger reef framework (as described in De Goeij et al. 2009, Alexander et al. 2015). Any epibionts present on the substratum were removed. Sponges were moved to 100-L running seawater aquaria (± 26°C; flow speed 3 L min^-1^) and kept under semi-transparent black plastic sheets to imitate *in situ* ambient light conditions of approximately 5–15 µmol photons m^−2^ s^−1^ photosynthetically active radiation (PAR) during daylight hours. Sponges were acclimatized at least 1 week prior to start of experiment.

Subsequently, tissue samples were collected for RNA extraction either before wounding (‘untreated’), at 2-h post-wounding and 12-h post-wounding. Wounding was done according to Alexander et al. (2015), by cutting a 1 cm^2^ piece of sponge tissue from each specimen with a sterilized surgical blade. Tissue samples (untreated, 2-h, and 12-h post-wounding; *n* = 3 per treatment) were taken from the immediate surroundings of wounded and regenerating tissues, out to a maximum of 0.5 cm from the site of initial wounding (see Alexander et al. 2015, carefully avoiding any attached coral rock, and removing this if present) and then flash frozen in liquid nitrogen, transported on dry ice to the Netherlands and stored at −80° until further processing. Samples were then sent on dry ice to Vertis Biotechnologie AG (Freising, Germany) for RNA extraction, library preparation and sequencing.

Two individual sponge samples had RNA extracted successfully for the 2-h post-wounding stage samples, and both a single sample for the untreated and 12-h post-wounding sample. Tissues were ground in liquid nitrogen and total RNA was isolated from samples using the TriFast RNA isolation kit (peqlab). The two 2-h post-wounding samples were combined in equimolar amounts, and for the untreated and 2-h samples ribosomal RNA molecules were depleted using the Ribo-Zero rRNA Removal Kit (Human, Mouse, Rat) (Illumina). For the 12-h sample poly-A selection was performed and RNA was fragmented with ultrasound (1 pulse of 15 s at 4°C). First strand cDNA synthesis was performed with N’ randomised primers, and Illumina Truseq adapters were used for second round PCR (barcodes: CAGATC (untreated), GGTAGC (2-h), and AGTTCC (12-h)). Normalization was performed, using denaturation and re-association of cDNA, followed by hydroxylapatite column clean up. Gel extraction was then used to select cDNA of sizes 400–700 bp for sequencing. This was sequenced on the Illumina MiSeq system machine with 2x300 bp read length. Sequencing was performed on a single run, with all three samples pooled. Demultiplexing was performed by the provider before sequences were uploaded to their server.

### Transcriptome assembly and filtering

The quality of the raw reads was assessed and visualized using FASTQC v. 0.10.0 (Andrews 2010). Using TrimGalore v. 0.2.6 (Krueger 2012) adapter, sequences and bases with low quality Phred scores (< 33) were trimmed and removed, and we applied a length filter to retain sequences of > 25 bases. Following this, high-quality reads were re-screened in FASTQC to ensure sample quality.

As the genome of *H. caerulea* is not available, four *de novo* assemblies for the species were produced. The three treatment libraries (untreated, 2-h, and 12-h post-wounding) were assembled individually, and a fourth complete ‘reference’ transcriptome for the species was assembled including all of the reads from three different treatments combined into a single assembly. All *de novo* assemblies were performed with Trinity 2013_08_14 (Grabherr *et al.* 2011) with a *k* mer size of 25 and all default parameters except for path reinforcement (50), and minimum transcript length of 200 base pairs (bp). Transcripts shorter than 200 bp were removed as shorter transcripts have been shown to exhibit poor similarity and retrieve few results in blast queries against sequences of proven homology (Riesgo et al. 2012). Assemblies are available to download in full from https://doi.org/10.6084/m9.figshare.5178217.

The four *de novo* assemblies were blasted against a subset of the NCBI *nr* database containing only proteins from Metazoa, using BLASTx (Altschul et al. 1997) with a cut-off *E*-value of 1*e*^-5^. Blast results served as a database for identification of genes found in the assemblies, and for those with different relative expression between treatments. In addition, given the large number of symbionts found within sponge tissues, the four *de novo* assemblies were blasted against all bacterial protein sequences in the NCBI *nr* database, using BLASTx with a cut-off *E*-value of 1*e*^-5^. Transcripts with hits only against the bacterial database (with no homology to metazoan proteins) were removed from subsequent analysis unless otherwise indicated.

### Transcriptome assessment, annotation and differential analysis

To assess transcriptome completeness, BUSCO (Simão et al. 2015) was run against only those contigs with a hit to a metazoan sequence in the *nr* database. The eukaryota_odb9 and metazoa_odb9 datasets were used for comparison, with all other default settings. We performed automated annotation using standalone BLASTx (Altschul *et al.* 1997) to the metazoan portion of the *nr* database, with a cut-off *E* value of 1*e*^-5^. Blast hit results of the four *de novo* assemblies served as basis to retrieve Gene Ontology (GO) terms using Blast2GO (Conesa *et al.* 2005) under the three overall categories: biological processes, molecular function, and cellular component, hierarchically organized into different levels. GO enrichment analyses using Fisher’s exact test were also performed in Blast2GO to test whether any of the GO terms appeared significantly over or under-represented in pairwise comparisons between the treatments (untreated, 2-h, and 12-h post-wounding). The significance threshold of the *p*-values for multiple comparisons was adjusted to a 0.05 false discovery rate (FDR) (Benjamini and Hochberg 1995). Automatic annotation was also performed using the KEGG-KAAS online server (http://www.genome.jp/tools/kaas/, Moriya et al. 2007, Kanehisa et al. 2016) using the blast strategy, the SBH method and a variety of eukaryote target species, including *A. queenslandica.*

Using the GO annotation results for each transcriptome, we obtained the frequency of GO terms in each treatment and calculated the relative increase for the untreated vs. 2-h and untreated vs. 12-h samples (Supplementary Files 1, 2). To visualize the GO terms that increased in each comparison we used REVIGO web server (Supek *et al.* 2011), graphically representing the results with the “treemap” R package. Size of the rectangles was adjusted to reflect the relative frequency. Further differential expression analysis was performed with the Trinity module (incorporating RSEM and edgeR, Grabherr et al. 2011) to map individual libraries with the total reference assembly. edgeR was run with a p-value cut off for FDR of 0.001, a dispersion value of 0.1 and min abs(log2(a/b)) change of 2 (therefore, minimally, 4-fold change). Both ‘as-isoform’ and ‘as-gene’ data are provided in Supplementary File 3 and 4.

### Manual annotation and investigation of target genes

A number of metazoan genes previously implicated in apoptosis, cell-cell interactions, cell proliferation, cell shedding in the gastrointestinal tissue, inflammatory processes, collagen production, and wound healing were selected based on prior appearances in the literature. Protein sequences of these genes with established homology were downloaded from the NCBI *nr* database, and used as queries to search a local database of our assemblies using tBLASTn (Altschul *et al.* 1997). Putatively identified hits within our assembly were then blasted individually against the *nr* database to confirm homology by reciprocal blast, and the presence of characteristic domains and residues was used for further corroboration when necessary. Initial searches were performed on the reference transcriptome, and when no sequences were found in this, we examined individual assemblies to confirm absence, or identify sequences that were not found in the overall resource.

## Results and Discussion

### Sequencing quality and depth

We obtained 15,749,700 reads for the untreated sample, 9,351,920 for the 2-h post-wounding sample, and 9,003,560 for the 12-h post-wounding sample (Table 1). Initial review with FastQC revealed the presence of low sequencing quality (Phred score > 33) bases at the ends of some reads, especially in the last 100 bp, and that some adaptors had not been removed in pre-processing. GC content was also variable, with 42.5%, 42% and 45.5% observed in untreated, 2-h, and 12-h post-wounding samples, respectively. This variability could suggest the presence of errors or bias in some of our reads before clean up, and we therefore were aggressive in our treatment of reads pre-assembly. Removal of adaptors and low quality bases with TrimGalore lead to between 15.4% and 21.5% of reads being removed completely (as they were less than 25 bases long) and a total of 27,807,356 reads remaining for all samples. Post-cleaning, all reads were of good quality, with an average GC% of 41%, 41%, and 44% in the three samples. While some over-represented sequences were observed in our reads, these were not excessively common, had no known blast hits and were almost entirely due to overly common *k*-mers at the start of reads, almost certainly due to known biases in Illumina hexamer binding (Hansen et al. 2010). Raw and trimmed reads have been uploaded to the SRA, with accession PRJNA371551 (NB currently embargoed, but will be released alongside publication).

**Table 1:**
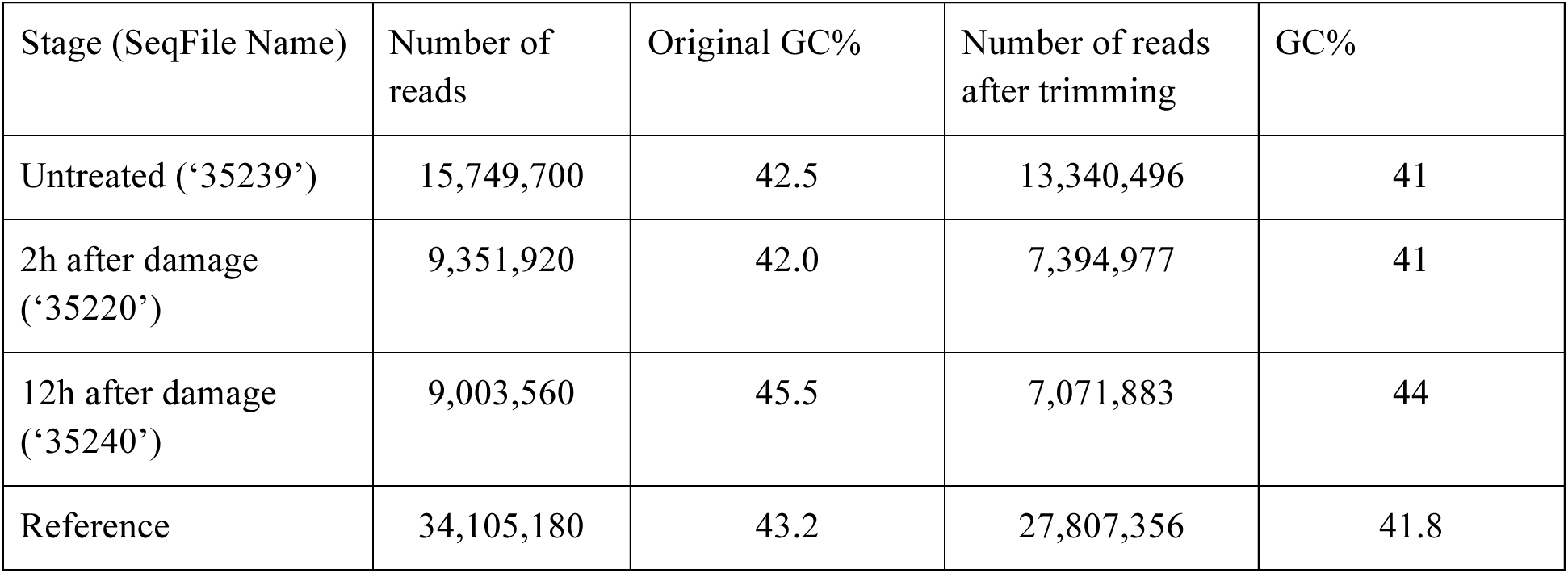
Basic statistics regarding sequencing and read quality.

### Transcriptome characterization: assembly and assessment

All four assemblies are of good to excellent sequencing depth, with in excess of 100 million assembled base pairs (bp) and with a high N50 (Table 2). However, the 12-h post-wounding sample shows less transcriptional diversity as assessed in raw total bp, albeit with better contiguity, higher average lengths and better annotations, relative to the other three assemblies. All assemblies contain a significant number of contigs greater than 1000 bp in length and N50s were always close or exceed 1,000 bp (minimum 836 bp, maximum 2032 bp). This is particularly useful as these longer contigs are more readily annotatable, as their longer length means that they span known coding domains and provide more information for both BLAST-based and manual annotation efforts. Resulting assemblies are available to download in full from https://doi.org/10.6084/m9.figshare.5178217.

**Table 2:** Basic statistics for assemblies.

| Stage | Number of transcripts | Number of Trinity 'genes' | Total bp in assembly | % G+C | N20 | N50 | Number of transcripts over 1000 bp | Transcripts > 1000 bp w/ blast hit | Transcripts > 1000 bp w/ GO term |
| --- | --- | --- | --- | --- | --- | --- | --- | --- | --- |
| Untreated | 282,239 | 154,516 | 203,062,338 | 42.08 | 2,114 | 836 | 54,706 | 24,487 | 2,265 |
| 2h after damage | 271,240 | 159,968 | 190,402,491 | 42.25 | 2,142 | 957 | 50,047 | 24,961 | 1,968 |
| 12h after damage | 113,254 | 69,503 | 105,485,222 | 43.22 | 3,771 | 1,688 | 30,000 | 22,608 | 2,655 |
| Reference | 515,624 | 169,711 | 600,590,960 | 43.18 | 3,998 | 2,032 | 189,770 | 138,309 | 2,413 |

Trinity is a splice-aware assembler, and groups similar contigs as “genes”. This process is not perfect, but can be an initial guide to the depth and transcriptional diversity of a transcriptome assembly. All of our individual assemblies possessed 1.6–1.8x as many contigs as inferred “genes”, indicating that some splice, allelic or other diversity in coding sequences was recovered by the Trinity assembler. However, our ‘reference’ assembly possessed approximately three times as many transcripts as inferred ‘genes’, and only slightly more inferred ‘genes’ than our normal sample on its own (15,159 Table 2). The fact that only slightly more ‘genes’ are recovered suggests that we have sampled almost the total transcriptional diversity of *H. caerulea.* The increase in transcript number has two possible, and not exclusive explanations (1) the allelic diversity between the three sponges assayed in the individual transcriptomes (as these were specimens from different individuals) or (2) the increased sequencing depth gained by combining all reads together, which allowed the assembler a more complete overview of the transcriptional diversity of the species, with less “gaps”, allowing more successful assembly.

Sponge transcriptomes and genomes can contain contamination from other sources, including bacterial symbionts, despite poly-A selection. To assay whether this was the case in our assembly, we blasted our total (reference) assembly against both the bacterial and metazoan portions of the *nr* database. Of the 515,624 transcripts within this dataset, 433,011 had no resemblance to bacterial sequences with a fairly relaxed cutoff value for assessment (*E* 10^-5^). The maximum bacterial contamination from bacteria in our dataset as assessed by this measure is therefore 16.02%. This figure will be a deliberate overestimate, as many metazoan sequences with well-conserved protein domains will possess blast hits to both bacterial and metazoan sequences at this *E-*value cut-off point. We therefore did not remove these sequences immediately, but assessed which contigs in our dataset also possessed putative homologues in the metazoan portion of the *nr* database. It is worth noting the presence of bacterial sequences, however, both to flag contamination, and also as they could be useful for future work. In particular, these sequences could be used in future work on the impact of trauma on the bacterial fauna present on *H. caerulea.*

*H. caerulea* has been described as both a low microbial abundance (LMA) and a high microbial (HMA) sponge (Maldonado et al. 2012; Alexander et al. 2014), as it lacks high bacterial abundance in TEM histological sections (pointing to LMA), but has all the tissue features of a HMA sponge (see Weisz et al. 2008). In all sponge tissues a certain percentage of microbial sequence is expected, and even though 16.02% seems higher than previous results from LMA sponges, which are usually between 5 and 8% (Riesgo et al. 2014), this figure generally corresponds with low abundance of bacteria. Changes in both overall bacterial diversity and in specific symbiotic species could be of interest to future studies, and we therefore provide an unaltered complete dataset here.

To assess the completeness of our dataset, we took only the 205,069 contigs in our reference transcriptome with a hit to the metazoan portion of the *nr* database, and assayed them for the presence for BUSCO sets of genes. These are lists of genes generally found in single copy in their respective clades, and are excellent means of assaying transcriptome and genome completeness. Of the 303 genes in the eukaryote BUSCO cassette, only 1.3% (4 genes) were missing from our reference transcriptome. 78% (237) were found to be represented by complete transcripts, and 20.5% were “fragmented” (on incompletely sequenced transcripts as inferred from length of known homologues, or split between multiple contigs). Similarly, of the 978 genes represented in the metazoan BUSCO group, 77.9% (762) were represented in their entirety in our transcriptome, with 14.7% partially represented, and only 7.4% of these genes not found. This evidence suggests that our transcriptome is of sufficient sequencing depth to recover almost all the expected complement of genes to be found in a poriferan species. By way of comparison, the *A. queenslandica* cDNA set is missing 1.6% of the eukaryote set (5 genes), and 4.9% (49) of the metazoan complement. Our dataset therefore compares favourably to that of the complete genome of a fully sequenced sponge species. To further confirm this finding, we annotated our contig database using both automated and manual methods.

### Automated Transcriptome Annotation

We performed automated annotation using standalone BLASTx (Altschul *et al.* 1997). Blast2GO (Conesa *et al.* 2005) was then used to map GO terms and provide further annotation to our dataset. Out of the 515,624 transcripts of the reference transcriptome, 205,069 (39.8%) retrieved a blast hit from the metazoan selection of the *nr* database, with approximately 67% of them longer than 1,000 bp (Table 1). Unsurprisingly, the species with the highest number of best blast hits was *A. queenslandica,* with 60,056, followed by the cnidarian *Nematostella vectensis* (12,567). Deuterostomes such as *Saccoglossus kowalevskii* (9,811), *Branchiostoma floridae* (9,720) and *Strongylocentrotus purpuratus* (6,781), previously noted for their slow rate of molecular evolution, are also well represented in best blast hit statistics. The most surprising entry (5th in the top ‘best hit’ species) is *Pantholops hodgsonii,* the Tibetan antelope, with 9,292 best blast hits. We can rule out sample contamination from that source, but the genomes of the nematode genus *Steinernema* (Dillman et al. 2015: Supplementary File 2 S2A) also show a remarkable affinity to that species, and this may indicate shared bacterial contamination or slow rates of molecular evolution.

In total, using Blast2GO, 148,474 GO terms at all levels of the GO hierarchy were mapped to 76,070 of our contigs. A low fraction of the total contigs, and about half of those with a GO term, 35,477 (7% of the total assembly), were fully “annotated” by Blast2GO (Table 1) at the default annotation rule cutoff (55). While this is a small fraction, poor mapping could be the result of the differences between sponge sequences and those generally present in the databases (InterProScan, etc) used for GO annotation. We should also note that as 30,060 genes were predicted in the complete *A. queenslandica* genome (Srivistava et al. 2010), this number (even with some allowance for transcript diversity) is not substantially different to that figure. Further, 9,181 enzyme codes were assigned to 8,431 sequences in our dataset, and this would represent the lion’s share of enzymes within a typical metazoan gene cassette.

When the three individual treatments making up our dataset were considered individually, between 16 and 31% of transcripts retrieved blast hit from the metazoan selection of the *nr* database, and only between 3.4 and 2.3% of the total transcripts were assigned GO terms (Table 1). Similarly to the reference transcriptome, approximately 10% of transcripts in all treatments were removed because they obtained hits against the bacteria selection of the *nr* database. In all cases, transcripts over 1,000 bp always retrieved more blast hits and GO terms than those shorter than 1,000 bp (Table 1).

Using the peptide sequences predicted by Trinity, our dataset was annotated by blast similarity and KEGG-KAAS, revealing the recovery of many important cellular and inter-cellular pathways. 128,764 of the 272,404 peptide sequences predicted by our pipeline showed some similarity to proteins in the KEGG database, and as a result almost all KEGG maps were well-recovered (Supplementary File 5).

To demonstrate the recovery of key pathways by our dataset, we investigated the Wnt family of secreted signalling peptides, which, among their many responsibilities, are well known for a role in regeneration, and have been previously catalogued in the related sponge *H. dujardini* (Borisenko et al. 2016). The complement of *H. caerulea* is shown in Fig 2. This figure is based on KEGG mapping of genes in this pathway in *H. caerulea,* along with the completed genome of *A. queenslandica*, and almost all genes that would be expected in a sponge dataset can be seen in our sample (green in Fig 2). Many genes in these pathways are not observed, but were similarly absent from the *A. queenslandica* KEGG map of these pathways, while only three genes (*c-jun, Rbx1* and *Wnt1 1*) were present in that genome but are not found here. This could be the result of true loss, sequence divergence, or through these genes not being transcribed at the time points sampled. The presence of the Wnt cascade of genes in all our samples indicates that these genes may be involved in both normal processes and in regeneration responses. The excellent recovery of the almost total componentry of these pathways further suggests that our sequencing is of sufficient depth to recover the lion’s share of the transcribed cassette of genes of *H. caerulea.*

**Fig 1.**
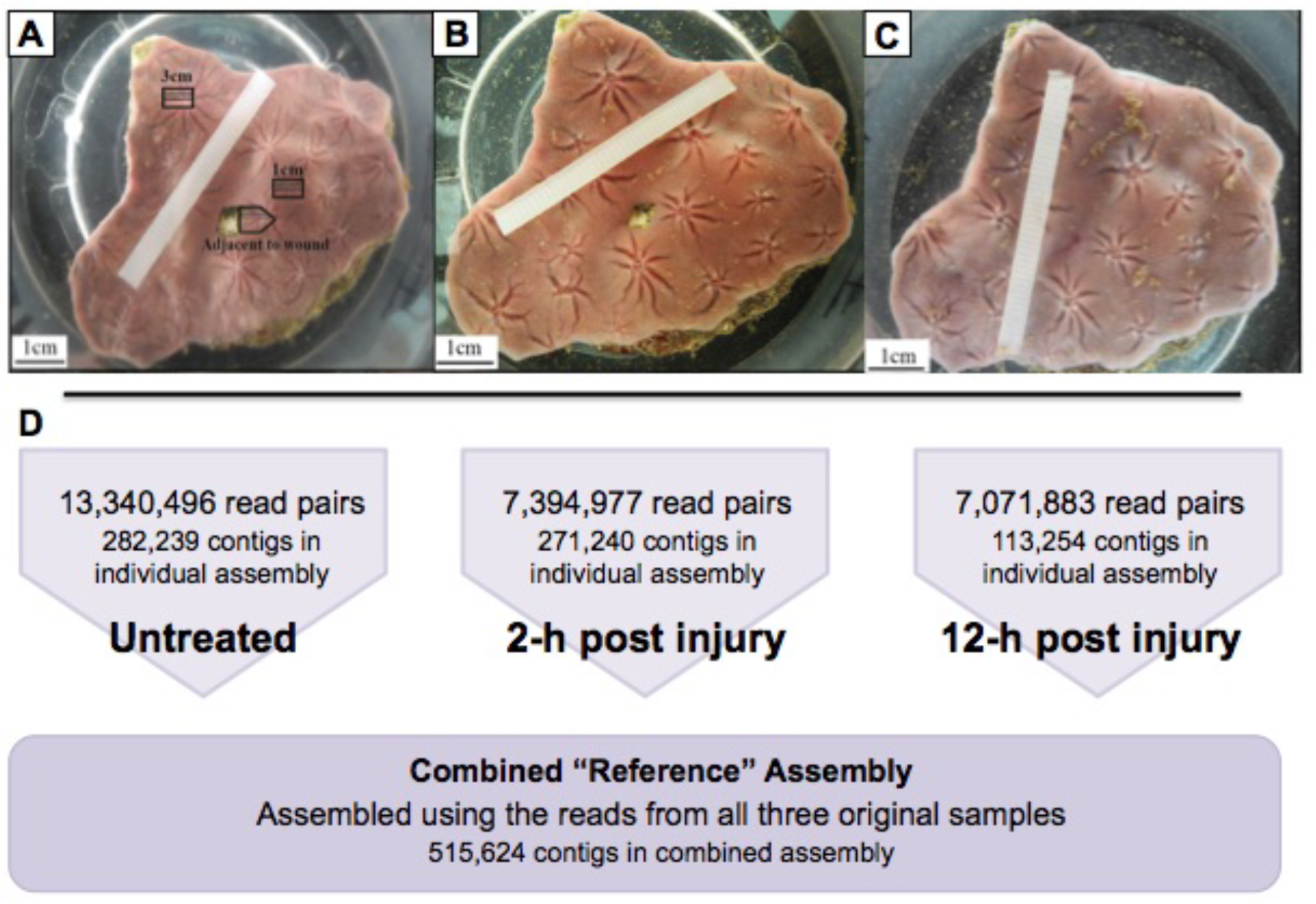
A) Image of a) *H. caerulea,* at the time of wounding, showing example of space “untreated” sample taken from, and adjacent space sampled for later analysis. Additional boxes show samples taken at set distance from wound in previous experimentation. B) image of *H. caerulea* one day after wounding, and C) after 6 days of recovery, note total regeneration. Images from Alexander et al 2015, doi: 10.7717/peerj.820, reused under the terms of the Creative Commons Attribution License. Below, D: Summary statistics, timed and reference transcriptome assemblies.

**Fig 2.**
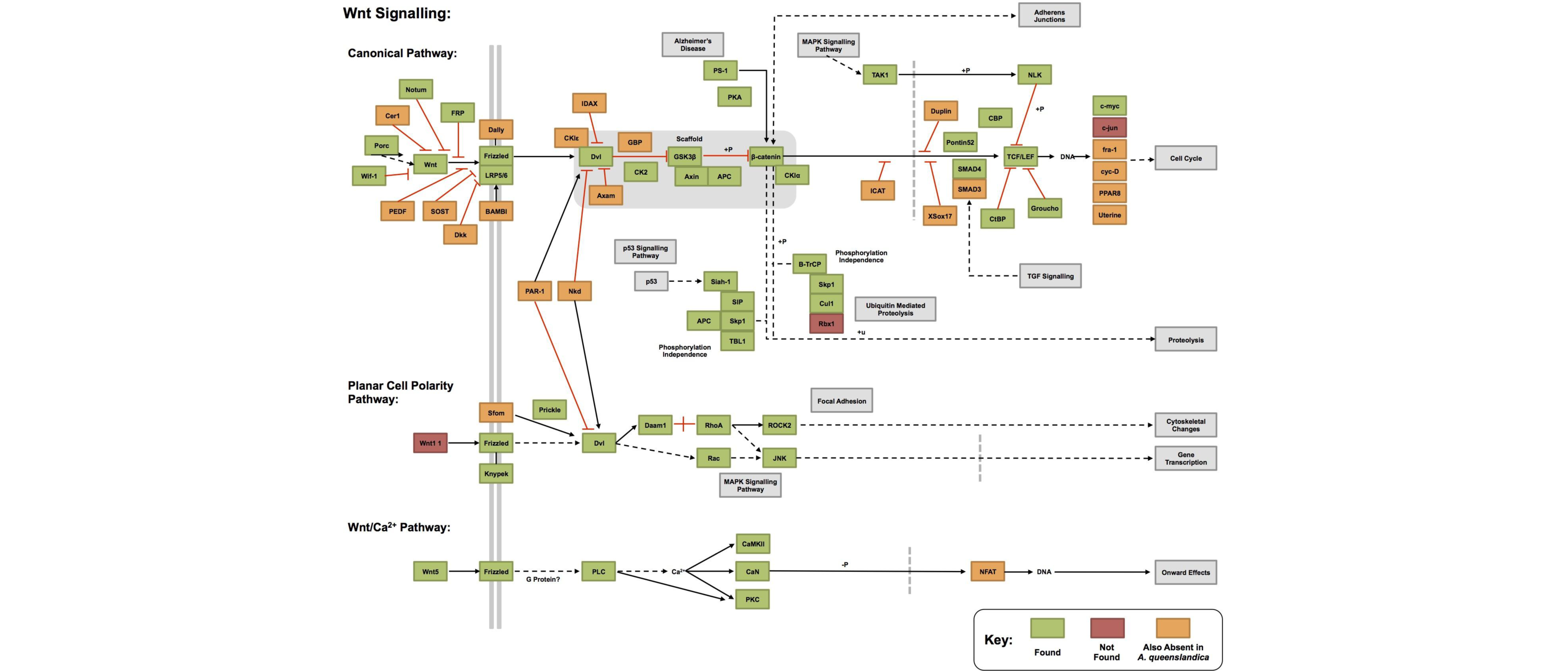
Recovery of canonical and non-canonical Wnt pathways in the Metazoan-only contig dataset described here, as annotated by the KEGG-KAAS Automatic Annotation Server Ver. 2.1 (Moriya et al. 2007, Kanehisa et al. 2016). The majority of all pathways examined are recovered in our resource. Some genes, marked in orange, are absent from our dataset, but may be absent from sponges more generally as they are coded as absent in the KEGG resource (normal sponge model: *A. queenslandica*). However, three genes appear to be absent from our dataset, but were previously identified in *A. queenslandica* (Srivistava et al. 2010). Assessment of genes as present is based on automated methods, and may be spurious if marked sequence divergence between genes in our dataset and orthologs in the KEGG database exist.

### Differential Gene Content of Timed Samples

The samples taken at each treatment point were normalised, and each was sequenced only a single occasion without replicates. The data presented here are therefore not suitable as the basis for quantitative analysis of gene expression levels. However, they provide a clear source of comparative information regarding gene presence, as this can be inferred directly from sequenced reads.

To gain an overall understanding of the changes in gene sequences present in our sample from time point to time point, we visualised the relative GO composition in the REVIGO web server (Supek *et al.* 2011 (Fig 3). The untreated vs. 2-h post-wounding sample shows a far more diverse range of high level GO terms than the untreated vs. 12-h sample, and in the 2-h sample these often possess a large number of child terms within their range. Many of the GO terms commonly found in the 2-h post injury sample (compared to the untreated one) are of apparent meaning. Contigs mapped to GO terms related to apoptotic processes are more common in that sample, along with contigs with GO terms related to response to biotic stimulus, embryo development (although these will overlap with development more generally), regulation of hormone levels and a variety of GO terms associated with structural components and metabolism (e.g., fluid transport, glucan metabolism, cellular metabolic compound salvage). These are all consistent with the observable processes of regeneration actively occurring at this stage. We also should note the presence of two GO terms that are upregulated in the 2-h sample - “entry into other organism involved in symbiotic interaction” and “dissemination or transmission of symbiont from host”. This specific process would be necessary for re-establishing the normal symbiotic complement of regenerated tissue, and aid the sponge in its survival, preventing other, less benign species from becoming established in this vulnerable period.

**Fig 3.**
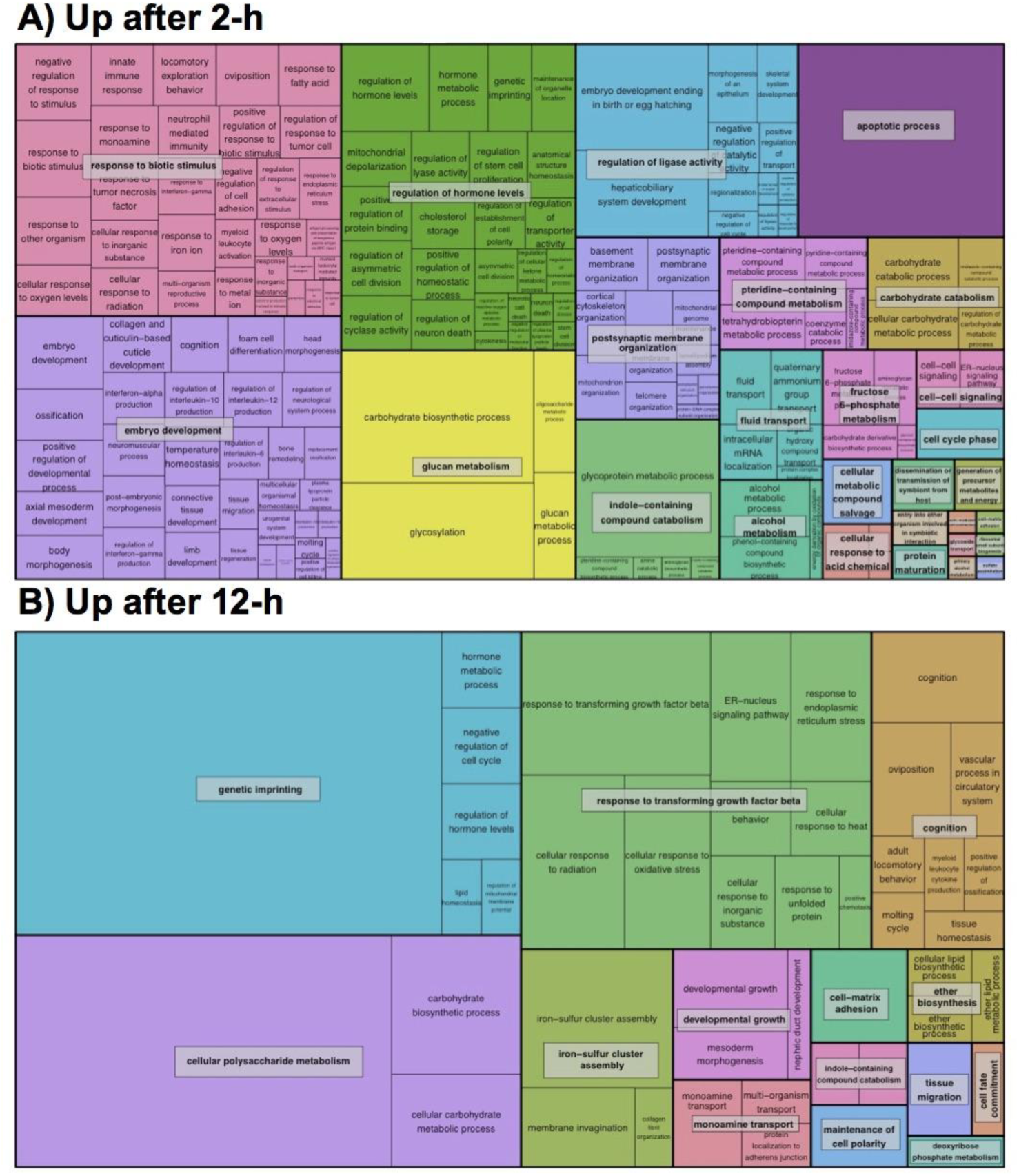
Regenerating *Halisarca caerulea* transcriptomic sample GO category representation, visualised using REVIGO (Supek et al. 2011). Note marked changes in GO category represented between these two time points, with little overlap in GO terminology between these times.

In contrast, comparison of 12-h GO term presence with that of untreated samples reveals fewer numbers of GO categories showing differences, and a very different representation of GO categories, with little to no overlap to the 2-h sample. The GO terms found to be most common compared to the untreated sample are those related to genetic imprinting, which may reflect cells taking up set roles within regenerated tissue. ‘Cellular polysaccharide metabolism’ and ‘response to transforming growth factor beta’ are the next-most different categories in representation relative to the untreated sample. This is consistent with cells that are taking up defined identities and establishing themselves in their roles after regeneration. Other categories upregulated in this 12-h sample and consistent with tissue re-modelling are terms such as “cell fate commitment”, “maintenance of cell polarity” and “cell-matrix adhesion”. The GO category “cognition” was also marked as more common, but while this seems unusual for a poriferan species, there are a number of gene families which are assigned to this category but are present in species without a CNS, including neurotransmitter synthesis pathways and their receptor genes. Sponges are known to possess many of these genes, even if their function in these species is not well understood (Sakarya et al. 2007; Riesgo et al. 2014; Francis et al. 2017). That these genes are involved in the late stages of a regenerative response may indicate a role in communication between cells, and deserves future scrutiny. The transcriptional difference between the 2-h and 12-h post-wounding samples point to a changing physiological status within the organism, transitioning from acute wound-healing towards regenerative processes. This is also clearly visible phenotypically, as after only a few hours the rims around the wound are already healed (Alexander et al. 2015). From 12-h post wounding onwards, it takes at least 6 days to fully recover in terms of cell proliferation (Alexander et al. 2015).

A more discrete understanding of the exact changes in gene expression was gained by looking at changes in expression of individual contigs between each sample. Within the Trinity software framework, we used RSEM and edgeR to perform a preliminary differential expression analysis, aimed at identifying possible novel target genes with roles in regenerative responses. We must stress the fact that no biological or technical replicates were performed, however, the results presented here will be of preliminary utility to future hypothesis-lead research.

Of our samples, the untreated and 2-h post-wounding samples were the most transcriptionally similar, as assessed by the Pearson correlation matrix of pairwise comparisons across all samples (Fig 4A, Supplementary File 4), (both “as isoforms” and “as genes”, in the latter case with contigs clustered automatically into groups, with single ‘genes’ representing multiple isoform data). The 12-h sample is markedly different in transcription levels for a variety of contigs (Fig 4B). This will at least in part be a consequence of the differing methods of library construction used for the 12-h timepoint in particular, as ribosomal RNA molecules were not depleted using the Ribo-Zero rRNA Removal Kit for this particular sample, as the sequencing provider did not deem this necessary.

**Fig 4.**
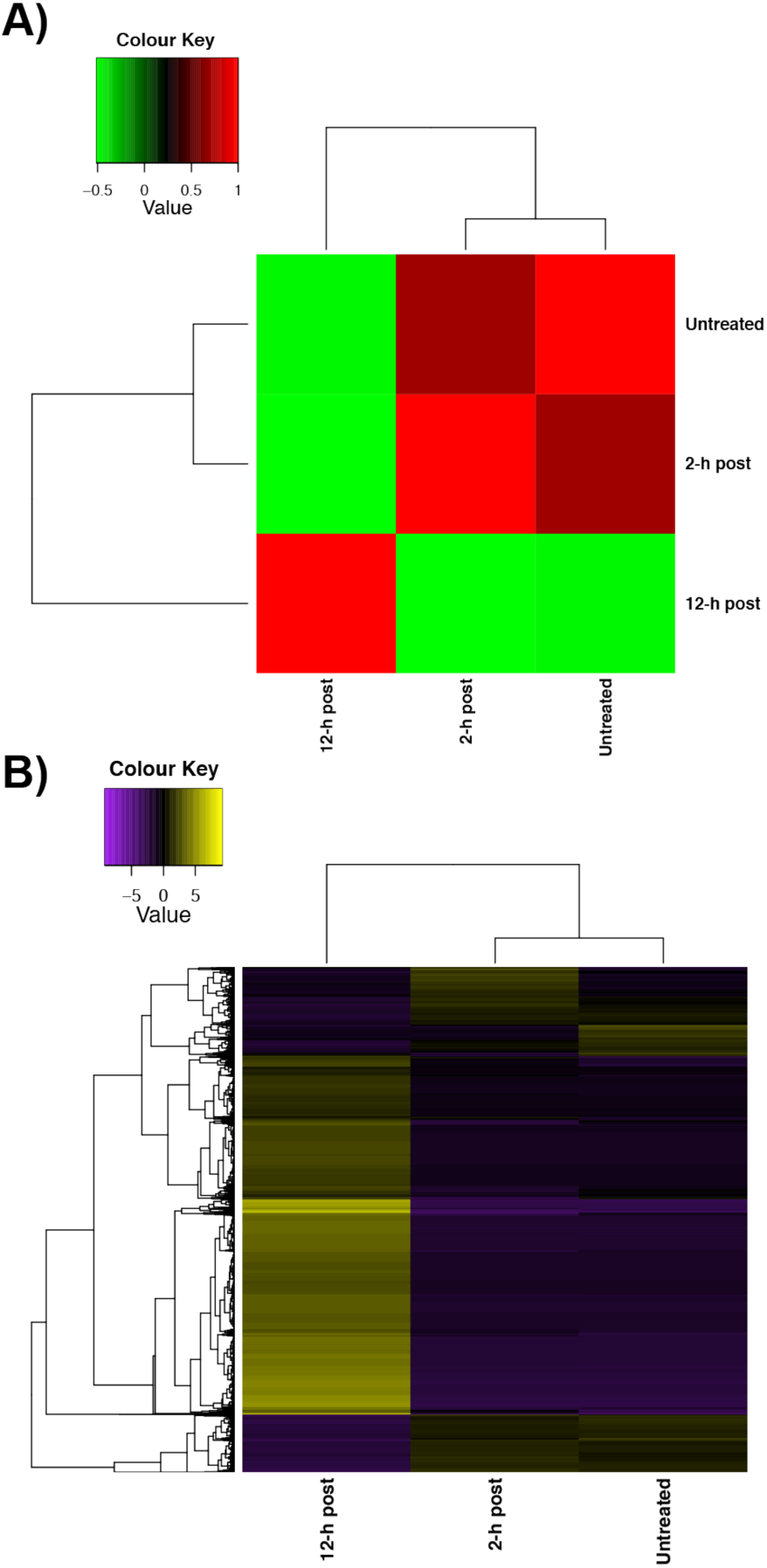
Cross-comparison of contig expression at the three time points examined in this work. This was performed by RSEM/edgeR within the Trinity framework, but should be taken as a guide to basic expression levels, particularly presence/absence, rather than a quantitative differential expression analysis due to a lack of replication. Note similarity of expression profile in untreated and 2-h samples, but marked differences in genes present at the 12-h stage, indicating marked transcriptomic remodelling. “As isoform” data shown here, but “as gene” information also available in Supplementary Files 3 and 4.

The impact of this, and the difference in transcribed cassette that is present in the 12-h sample can be easily seen in Fig 4B, where a large number of transcripts (in yellow) are markedly more prevalent in this dataset than in the other two samples - these transcripts are not observed, or observed at very low expression levels, in either the untreated or 2-h post-wounding sample. While we re-emphasise that our dataset was not sequenced in a manner that makes quantitative comparison valid, in a qualitative framework, we examined the 10 genes with the highest and lowest relative expression in each of our samples (Table 3), in order to determine whether any particularly noteworthy changes in gene presence/absence could be gained from our data, and in particular to determine whether these highly visible contigs in the 12-h sample represented real *H. caerulea* genes, or potential ribosomal contamination. Note that in a small number of cases, ‘upregulation’ of one isoform happens concurrently with ‘downregulation’ of another, and this may result in no change in overall gene expression (genes indicated with an asterisk). All of the “uncharacterised” genes also have homologues in other species, and are therefore evolutionarily conserved. Future work examining these genes, and testing whether they may be involved in regenerative responses more generally across the Metazoa would be well worthwhile.

**Table 3:**
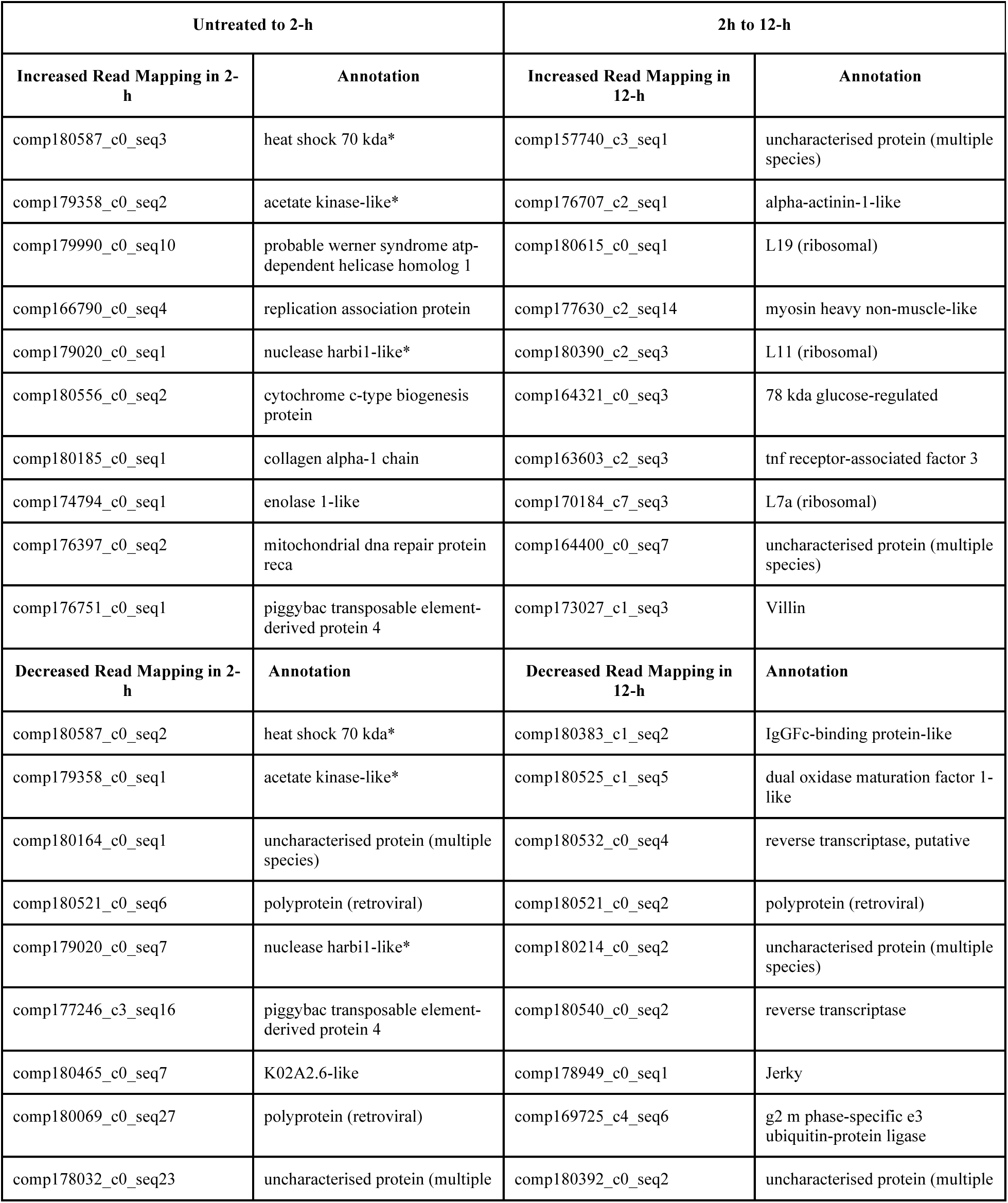
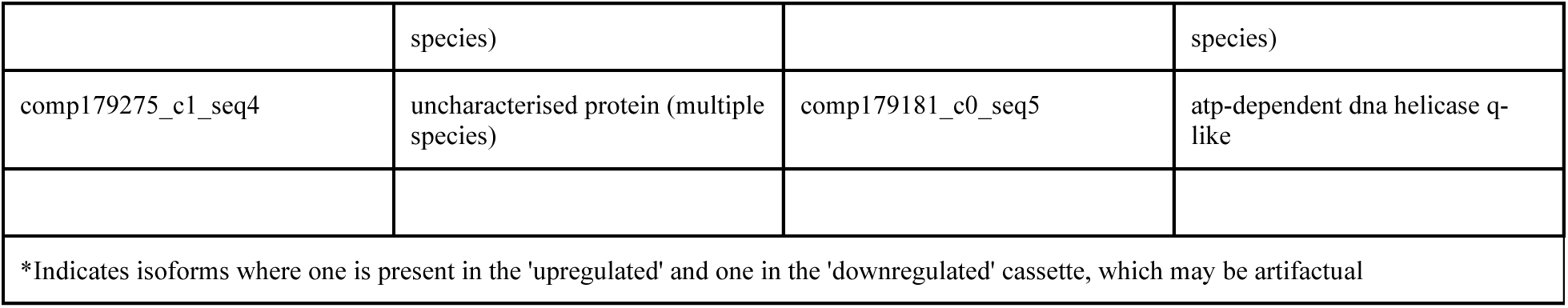
Top 10 biggest changes in read mapping to contigs, up and down, between our untreated and 2-h samples, and between our 2-h and 12-h samples. Annotation based on BLASTx identity.

Some genes on this list are of particular interest due to their likely role in regeneration: *collagen alpha-1 chain* is present in the list of genes more commonly present in the 2-h sample, and *alpha-actinin-1-like* and *villin* in the 12-h sample. Alexander and colleagues (2015; Fig 2) describe the formation of collagen (including collagen ducts) around and towards the wound area within 2 days after wound infliction. Collagen ducts are hypothesized to be used in (stem) cell transport towards the wound, to initiate cell proliferation at later stages of regeneration (6h–6d) after sealing the wound (early regeneration 0–6h after wound infliction, Alexander et al. 2015). In most animals, *collagen alpha-1 chain* appears as a component of the extracellular matrix (ECM) (Gelse et al 2003), while *alpha-actinin-1-like* and *villin* are usually found in association with actin filaments (e.g. Bretscher & Weber 1980). In a previous study of the cytology during wound healing in *H. caerulea*, a high abundance of collagen fibers were detected by picrosirius red staining in the regenerative wound tissue compared to areas of tissue away from the wound (Alexander et al. 2015). In addition, during regeneration in *H. dujardini*, the development of the ECM in the wounded area occurs approximately 12 hrs after injury (Borisenko et al. 2015), but in *H. caerulea*, this process could have proceeded faster. In this sense, while most genes of the ECM and basement membrane are expressed during the three timepoints, it is important to note that perlecan, an essential gene for basement membrane formation, is only expressed in the 12-h sample (Fig 5A). In addition, more transcripts with similarity to *collagen and calcium-binding egf domain-containing protein 1-like*, which is generally associated to endothelial cell migration and ECM formation (Hohenester and Engel 2002), appear in the sample collected 2 hrs after injury (Fig 5B). The increased read mapping of tnf receptor-associated factor 3 in the 12-h sample also provides an excellent novel target as a potential mediator of regeneration responses. Genes mapped to in lesser quantities in our analysis include reverse transcriptases and retroviral sequences. Why these would be transcribed in larger quantities in un-injured samples is unclear, but may be related to differences in “germ-line” composition - if most totipotent cells are involved in regeneration rather than reproductive division, these may be less well transcribed in our regenerating samples.

**Fig 5.**
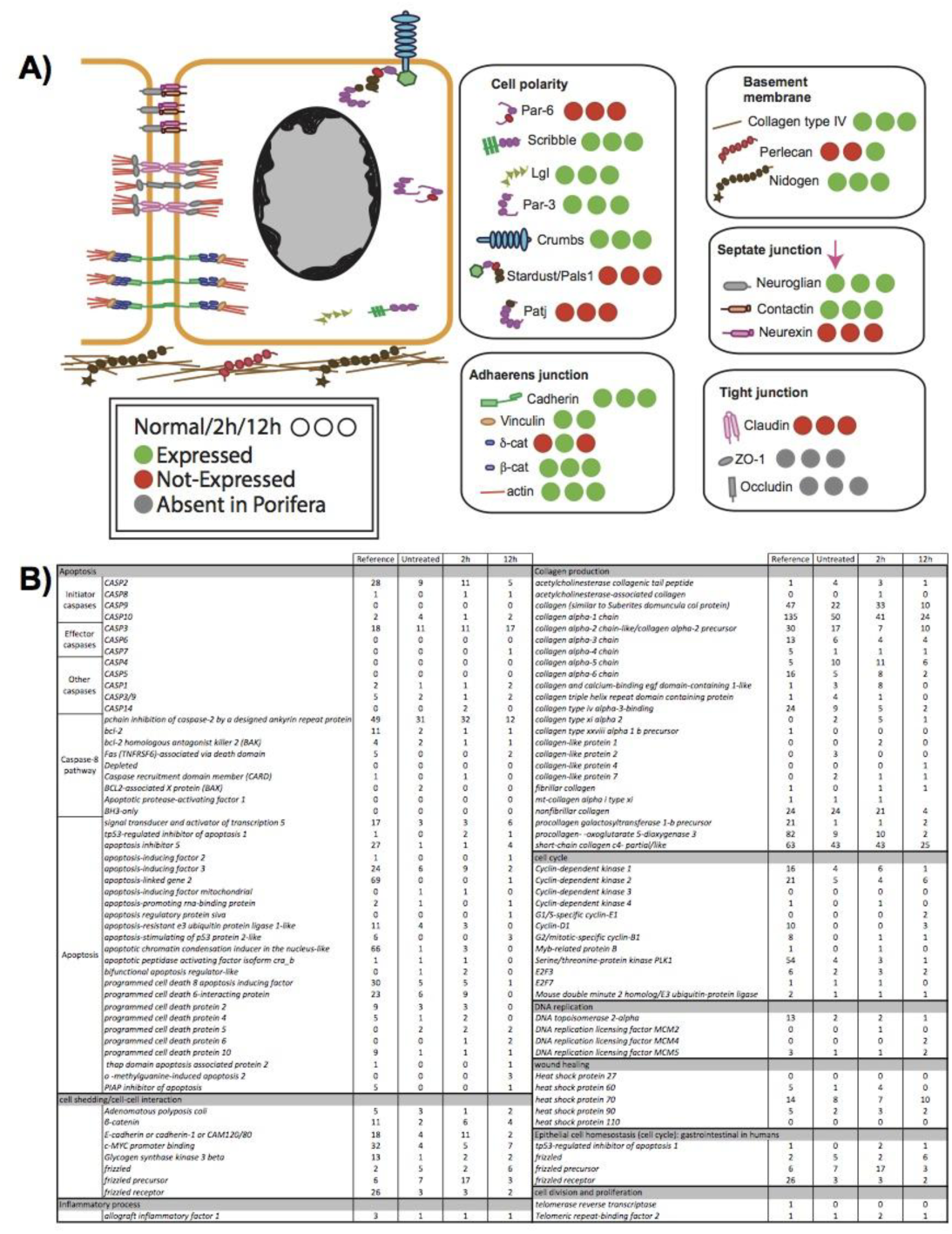
Genes involved in a variety of common cell cycle, death and signalling pathways, and their presence in the transcriptome of *H. caerulea* at the stages studied in this manuscript. A) shows the location of these diagrammatically, indicating in particular the location and presence of structural elements of sponge cells. B) indicates a further variety of these in tabular form, expanding further on this data, with full gene annotations available in Supplementary File 6.

Along with genes involved in regeneration, we noted the increased presence of a number of ribosomal protein genes in our 12-h sample relative to our 2-h time point. These are however ribosomal proteins, and not rRNA sequences. Upregulation of ribosomes may well be part of the regeneration response, as protein synthesis will be required for a variety of processes. We therefore believe that while the 12-h sample is perhaps partially skewed by an increased presence of ribosomal sequences, as noted above, due to the ribosomal component of this sample not being depleted before sequencing, this should not have too much of an impact on the utility of this time point. Further, this sample was also markedly different in transcribed content to the untreated and 2-h post-wounding samples, containing a large number of contigs not observed in our other two samples (Fig 5B). Only a small number of these contigs will be ribosomal, while the substantial non-ribosomal component of this data will be from other genes contributory to regeneration responses within this timed sample.

### Manually identified target genes

To confirm the presence of homologues of genes known to moderate regeneration in other species in our dataset we manually searched for and identified a variety of target genes in our transcriptomes. This allowed us to further verify the depth of our resource, initially assay for the potential role of specific in regeneration in the Porifera, and begin to build a picture of the sponge regenerative cassette for later hypothesis-lead work. We examined a variety of pathways in this work, blasting initially our combined reference transcriptome, which possessed the longest sequences for our target genes, with sequences of known orthologues of our genes of interest. We then secondarily retrieved orthologues from our untreated, 2-h and 12-h assemblies using their now-known *H. caerulea* sequences, alongside the sequences of our known orthologues from other species. Often we found several variants of the same sequence, representing possible allelic variation, splice differences or assembly artifacts within our samples. Given that our samples were drawn from different individuals, which were pooled in our combined “reference” transcriptome, this variation is not surprising, and the recovery of variation could be useful for further studies of specific gene families. The names of all these contigs and their annotation can be found in Supplementary File 6.

Given that regeneration must necessitate the re-establishment of tissue connections and boundaries, a variety of genes involved in junction formation and regulation were studied, (Fig 5A). This figure shows the location of these proteins in reforming sponge tissue, along with their presence/absence in our transcriptomic resource. Many genes are universally present in both our untreated and regenerating samples, and therefore cannot be linked with a regeneration response. However, *δ-catenin* and *perlecan* are only present in 2-h and 12-h post-injury samples respectively, and therefore may play a more distinct role in mediating regenerative responses at these times. While *δ-catenin* is involved in cell-cell adhesion processes, *perlecan* is the major proteoglycan in the basement membrane (Hohenester and Engel 2002). *Perlecan* has been previously found in several demosponges, calcareous and homoscleromorph sponges (Riesgo et al. 2014), although it is still reported as absent from sponges because it is not contained in the genomes of *A. queenslandica* and *Oscarella carmela* (Fidler et al. 2017). Collagen production is a well understood part of the wound healing response (e.g., Borisenko et al. 2015; Alexander et al. 2016), and we therefore looked for collagen components and related genes in detail. We found a plethora of collagen chain genes and related proteins, and these were found in a diversity of possible isoforms. As noted above, no particular pattern was noted in the amount of contig diversity between untreated and regenerating samples, but one gene (*collagen alpha-1 chain)* was noted as being upregulated in our 12-h dataset by our preliminary differential expression analysis, and it may be that the differential expression of certain isoforms occurs in regenerating samples, but at present we cannot comment on this in detail with the data at hand. Similarly, several *spongin short chain collagen* genes were found to be differentially present in our transcriptome samples (Table 4). Although some authors consider them to be homologous to *collagen type IV* (Aouacheria et al.2006; Fidler et al. 2017), they are indeed different molecules that co-occur with *collagen type IV* in all sponges, including *H. caerulea* (comp178545_c1 in this dataset, with numerous isoforms present).

**Table 4:** *Spongin short chain collagen* genes and their relative prevalence in our untreated and regenerating samples

| Trinity 'Gene' | # annotated isoforms/splice variants | Over-represented, Untreated | Over-represented, 2h | Over-represented, 12h |
| --- | --- | --- | --- | --- |
| comp150981_c0 | 1 |  |  |  |
| comp153402_c0 | 1 |  |  |  |
| comp162348_c0 | 10 |  |  |  |
| comp163102_c0 | 2 |  |  |  |
| comp165291_c2 | 1 |  |  |  |
| comp165464_c1 | 1 |  |  |  |
| comp167536_c2 | 2 | X (isoform 1) |  | X (isoform 1,2) |
| comp171120_c0 | 7 |  |  | X (isoform 6) |
| comp175460_c0 | 3 |  |  |  |
| comp176046_c0 | 1 |  |  |  |
| comp177461_c0 | 14 |  |  |  |
| comp177772_c2 | 1 |  |  |  |
| comp177821_c0 | 10 |  |  |  |
| comp178550_c0 | 2 |  |  |  |
| comp179613_c0 | 7 | X (isoform 17) |  |  |

We examined apoptosis-linked genes in particular, given their key role in tissue remodelling in regenerative responses, and apoptosis’ likely role in *H. caerulea* regeneration. Of the caspase genes, which are vital for programmed cell death and inflammation responses (Shalini et al. 2015) we found multiple sequences for the *CASP2* (initiator caspase) gene, and at least one sequence for other CASP gene members, with the exception of *CASP4, CASP5, CASP6* and *CASP9* (Fig 5B). A vast diversity of other genes linked to apoptotic responses were also studied, and these can also be seen in that figure. The presence of so many genes from this vital cellular cassette in our sample further confirms the depth of our transcriptomic resource. Further regeneration-related categories of genes we examined included cell shedding and inflammatory process genes (which are known to be specifically important in *H. caerulea* regeneration), cell cycle-related genes, heat shock proteins, and those related to cell division and proliferation. As with collagen and CASP family genes, these did not show any particular increase in diversity between untreated and regenerating samples.

### Utility of these resources for study of <u>Halisarca caerulea</u> regeneration

The transcriptomic resources described above were conceived as being a starting point for understanding the molecular elements of regeneration in *H. caerulea* in particular, and for finding the extent to which the molecular underpinnings of that response are similar to those found more widely in the Metazoa. Evidence from BUSCO, our recovery of key signalling pathways and our general recovery of a diverse range of gene sequences in our manual annotation attempts all suggests that we have succeeded in creating a deep reference transcriptome for this species. The far larger number of genes annotatable in our reference transcriptome, which in almost all cases is more than the sum of the individual transcriptomes, reinforces the power of increased sequencing depth in allowing complete recovery of transcripts, simplifying subsequent annotation. However, as a result of the non-quantitative sequencing procedure used in the construction of libraries we cannot comment conclusively on the relative expression of different contigs. Nevertheless, the preliminary data presented here can be highly useful to future, hypothesis-driven, research.

## Conclusion

Sponges have much to teach us about the ancestral regenerative cassette of the Metazoa, and may also hold clues for the promotion of healing and regeneration in a medical context. To date, however, our investigations of sponge regeneration have been largely limited to histological, cell dynamic and gross morphological studies, limiting our ability to cross-compare sponge regeneration more broadly with that of other Phyla at the molecular level. The advent of next-generation sequencing allows us to rapidly gain this understanding, and here we have described several resources that not only allow the first insights into the pathways regulating regenerative processes in the Porifera, but will also stand as excellent resources for continuing efforts in that clade. The reference transcriptome described here in particular will be a bedrock for future comparative work, while the conserved pathways noted as playing a part in this process are well recovered and show signs of differential expression between our samples. These initial findings will need to be confirmed and extended further by future work, but with this resource in place, our understanding will have a firm basis from which to proceed.

## Acknowledgements

The authors thank the members of their laboratories for many helpful discussions and all their support in preparing this manuscript. We thank the reviewers and editors of this manuscript for their suggestions and time, and CARMABI Foundation staff (Curaçao, Netherlands Antilles) for their help with collection and sample preparation.

## Supplementary Files

Supplementary File 1: GO annotations, all genes in reference transcriptome (.annot)

Supplementary File 2: GO categories upregulated, 2h, 12h samples, several files including Fishers’ Exact Test and REVIGO results (.zip)

Supplementary File 3: Genes/isoforms with significant differences in representation between samples when mapped, upregulation amount, annotation (.zip)

Supplementary File 4: Full results, comparison of expression, numerous files including volcano plots, full lists of genes up and down regulated (.zip)

Supplementary File 5: KEGG annotations, reference transcriptome (.txt, zip) Supplementary File 6: Blast annotations for all contigs in reference transcriptome (.txt, zip)

